# Online Forays for Macrofungi: Expediting the Rate of Documenting Biodiversity

**DOI:** 10.1101/2022.05.24.493314

**Authors:** Stephen D. Russell

## Abstract

A citizen-science oriented mushroom collecting event was organized online and held across the state of Indiana from October 23-29, 2017. Utilizing iNaturalist as the observational reporting platform, 30 members of the Hoosier Mushroom Society made over 1,300 observations of macrofungi during the week of the event. More importantly, the associated specimens were saved for more than 800 of the observations. Over 400 of the specimens were retained and received DNA barcodes, resulting in novel genotypes for 73 putative species being added to public databases. A full set of administrator and participant protocols has been compiled to aid others in conducting this style of biodiversity research for macrofungi. Methods include volunteer recruitment, participant protocols, specimen shipment, triage, and storage. Data management is also a key administrative aspect. Methodologies for integrating iNaturalist observational data with DNA sequence data and herbarium collection information are outlined for the MycoMap platform. Traditional mushroom collecting events have a number of limitations when compared to the online foray model. These advantages are discussed and key considerations for hosting online forays are examined.

## Introduction

Citizen science projects hold the promise of significantly enhancing the rate that biodiversity can be documented, particularly for macrofungi. In North America, large networks of local mycological societies and mushroom enthusiasts stretch across the continent, along with tens of thousands of individuals who form mushroom-related communities on social media (Sheehan 2017). Recent initiatives like the Fungal Biodiversity Survey (www.fundis.org) show great potential in engaging citizen scientists to photograph, document, voucher, and genetically sequence specimens of macrofungi across a broad scale. New methods for engaging these populations has the potential to open a previously untapped spigot of scientifically valuable data. Fully engaging citizen scientists could allow for significant gains to be made towards the understanding of North American macrofungal biodiversity, all while being achieved in much shorter timeframes than has been possible historically.

Most of the biodiversity of fungi remains unknown, despite their vital role in ecosystem function and human affairs. The most widely accepted estimates of total fungal diversity range from 1.5 – 5.1 million species, yet less than 10% of the species have been described in the literature (Blackwell 2011, Hawksworth and Rossman 1997, Hawksworth and Luecking 2017). Over the last 40 years, the rate of new species descriptions has remained consistent at around 1,300 species per year (Hawksworth and Luecking 2017) Assuming this rate is maintained, it will take well over a millennia to document the extant biodiversity. The current methodologies for cataloging fungal biodiversity are wholly insufficient when compared to the scope of the work that is believed to remain. During this period of an unprecedented loss of global biodiversity (Pimm et. al 2014), fundamentally new models for documenting fungi are required if many of these species are to be documented before becoming extinct.

Assessing the biodiversity of macrofungi (fungi that produce a macroscopic fruiting body) in a modern context is dependent upon vouchered specimens, typically with associated DNA barcode data (Bruns and Beug 2012, Korf 2005, Truong et al. 2017). Accurate identification of specimens in the field is often hampered by the significant macromorphological variation within a single genotypic species, or alternatively, convergent morphologic characters between separate genotypic species (Hawksworth and Lücking 2017; Korhonen, Seelan, and Miettinen 2018). DNA barcoding has been shown to be a robust and cost-effective means of identifying species of macrofungi (Schoch et al. 2012, Seifert 2009, Yahr, Schoch, and Dentinger 2016), yet despite widespread adoption in the professional scientific community, few citizen scientists have access to this important tool for their biodiversity projects.

Starting in 2015, members of the Hoosier Mushroom Society began work on a statewide survey of all the macrofungi that occur in Indiana. The ultimate goal of the endeavor is to create a web-based, open-access mycoflora for the state. This online compendium would include photographs, distribution maps, seasonality profiles, identification keys, and descriptions, all back-boned by vouchered specimens and DNA data for each species that is circumscribed. The state of Indiana covers a large geographic area; it can take well over 5 hours to drive from the northern part to the southern part of the state. While many individuals would like to contribute to the statewide survey, it is difficult to hold regular collecting events that can be attended by most members. Moreover, only a few members accounted for the large majority of specimens that were being preserved in the state. To give more citizen scientists the chance to make a meaningful contribution, survey organizers looked for an alternative model that would allow for more participation by the membership, especially from areas of the state that are under-documented.

### Project Goals

1.) Fully engage the citizen science community by reducing the barriers to participation.

2.) Train participants in making scientifically valuable collections of macrofungi.

3.) Maximize the number of valuable collections that can be DNA barcoded and permanently housed in local herbaria.

4.) Return results to survey participants.

5.) Develop a baseline of macrofungal biodiversity data for the state.

6.) Create a model project to capture biodiversity data that can be replicated by other organizations.

## Methods

A statewide, iNaturalist.org-based, online biodiversity survey for macrofungi was held on October 23– 29, 2017. This event was open to all members of the public who were willing to document mushrooms utilizing the iNaturalist mobile application and collect/dry the physical specimens of the mushrooms they encountered. This event is believed to be the first statewide, online mushroom foray known to have occurred. A full description of the methods employed for these events can be found below.

### Foray organizer preparation

#### Creating an iNaturalist project

In 2018, iNaturalist introduced two new project types – Collection and Umbrella (Seltzer 2018). These new formats broadly aggregate all iNaturalist observations from the website that meet specified parameters into a single project, whether they were observed by a participant of an independent effort or not. The older project type, a “traditional” iNaturalist project was utilized for the online foray. The “traditional” project type allowed individual observations to be included or excluded from the online project, allowing a project administrator to accurately monitor the success of their individualized effort, relative to all other iNaturalist observations being generated at the time. The “traditional” projects also gave project administrators options to require that observations have specific custom fields filled out by the participants as a prerequisite for addition to the project. These custom fields are called “observational fields” on iNaturalist.

The title for the iNaturalist project was “HMS Online Fall Foray 2017” (https://www.inaturalist.org/projects/hms-online-fall-foray-2017). The parameters that were utilized to set-up the project included an “open” membership model, an “anyone” preferred submission model, a location requirement of “Indiana,” and observational rules that limit submissions to the dates of the event. Finally, three “observational fields” were included in the project setup. They include “Voucher Specimen Taken,” which is a required field for submission to the project, as well as “Voucher number(s)” and “Collector’s name” which are not required to include an observation in the project.

#### Website

An informational page on the Hoosier Mushroom Society website was created to facilitate information to participants (Ex. - https://web.archive.org/web/20200926182627/http://hoosiermushrooms.org/index.php?/hms-online-fall-foray). This page included the dates for the event, protocols for participation, informational videos, and a compilation of frequently asked questions about the event. This website served as the primary informational portal to participants.

#### Volunteer recruitment

The majority of the recruitment efforts were centered around members of the Hoosier Mushroom Society (http://www.hoosiermushrooms.org) and members of the Indiana Mushrooms Facebook group (https://www.facebook.com/groups/indianamushrooms/). An email outlining the event and how to participate was sent to 1,522 individuals who had subscribed to the Hoosier Mushroom Society email list. These messages were sent in early October of 2017. Secondarily, leading up to the event, a series of posts were made to the Indiana Mushroom Facebook group that promoted the event and protocols. These posts linked to the informational page of the Hoosier Mushroom Society website and outlined potential prizes that would be given away for various levels of participation. Social media posts were also made to remind individuals to order field data slips, which was the lone pre-foray requirement for individuals who wished to participate.

#### Field data slips

Field data slips were utilized for this events (Figure 1). These slips have predefined sections where participants can take notes of the most important macroscopic features and environmental metadata that is significant when examining a macrofungal collection. They also have a ruler along the edges for scale. Most importantly, these notepads of 50 individual slips feature unique, consecutive numbers that are rendered in triplicate on each notepad. These numbers help to link the specimens that are collected with the iNaturalist observation numbers and help participants with organization during the process of drying the specimens. An order form was set-up on the Hoosier Mushroom Society website where participants could order slips before the event at no cost to them. Mailing the notepads cost between $1.00 and $2.00 per participant, and these shipping costs were covered by the event organizers.

#### iNaturalist mobile app

Participants were encouraged to download the iNaturalist mobile app and to become acquainted with it before the start of the foray. The mobile app is available on both Android (https://play.google.com/store/apps/details?id=org.inaturalist.android) or IOS (https://itunes.apple.com/us/app/inaturalist/id421397028?mt=8), so most participants with a smartphone should be able to participate. One key element that was emphasized to users was to join the foray project before they attempted to make their first uploads. This can be accomplished in the mobile application or on the iNaturalist website. Upon joining the project, mobile users have quick access to the project when creating new observational reports. Adding each observation to the project is critical, as that is a key trigger to ensure that the requested fields (Voucher Specimen Taken, Voucher number(s), and/or Collector’s name) are filled with the observation. Users who were not able to access the mobile app could upload their observational reports through the iNaturalist website.

**Figure 1.**
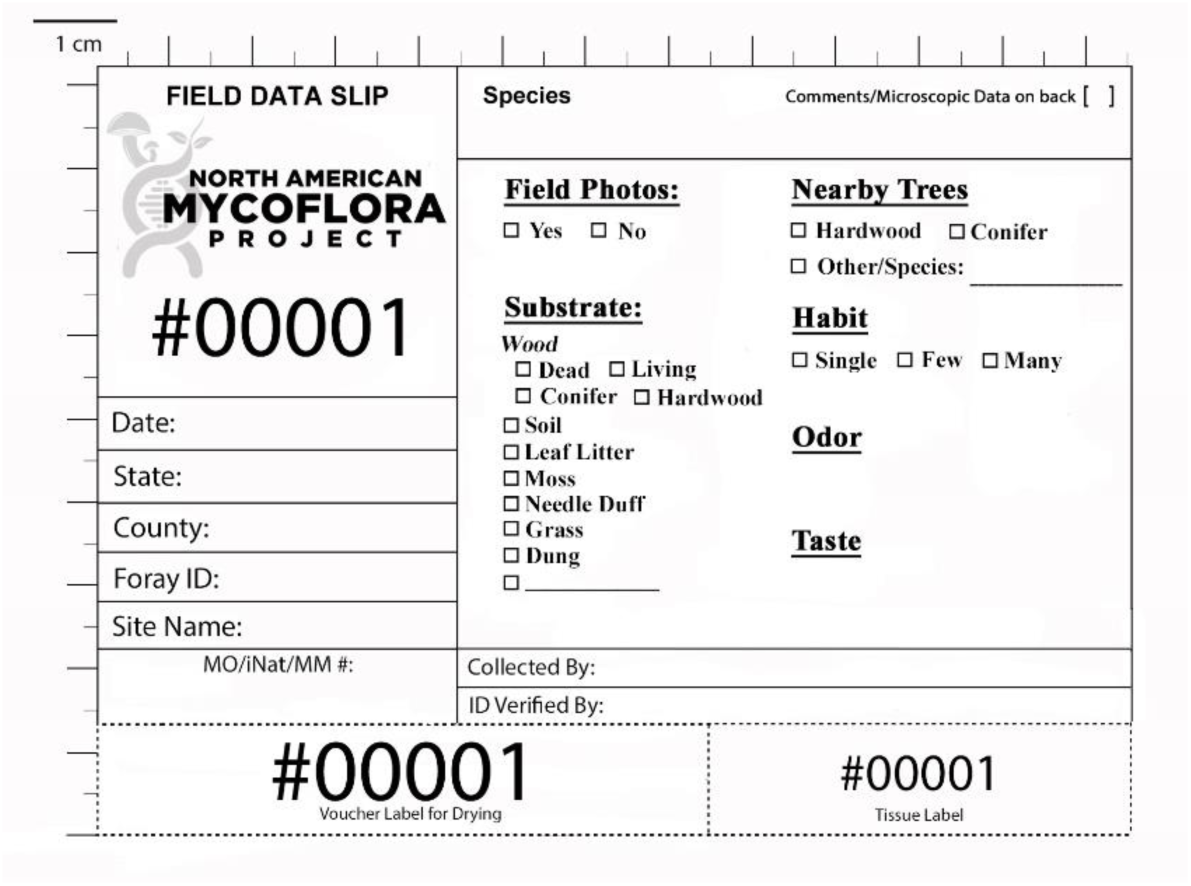
Field data slip. An example of a field data slip utilized for these events.

### During foray week – Participant protocols

The exact instructions given to foray participants are recreated below. An attempt was made to streamline the requirements for participants as much as possible, while retaining the most important metadata needed for individual collections.

**1) Create new observations of mushrooms you encounter**. This can be done through the iNaturalist mobile app or web interface. With each new observation, be sure to select the project for your event and whether you collected the specimen. The mobile app uploads the photos to online “observational” reports.

**a) Take multiple photos of the mushrooms with your cell phone or camera**. Take a nice image near ground level from the side, as well as an image of the top, the stem, and the spore bearing surface (the gills or pores on the underside of the cap).

**b) If you intend to save the specimen, take an image of the field data slip with the specimen**. This can be done in the field or back at home. Be sure the scale on the side of the slip is visible next to the specimen.

**c) Enter the field data slip number into the “Voucher number(s)” field in the mobile app**.

**2) Collect the specimen. Store the entire field slip (or detach the portion with the number) and place it with the specimen**. Fill out your field slip completely with the requested information before you dry your specimens.

**2) Back at home, dry the specimens with a dehydrator or fan**. Use the duplicate number at the bottom of the voucher slip to organize collections as they are being dried (Figure 2). Tape this number near the specimen. Once they are cracker dry (usually 1-2 days) put the voucher slip and the specimen in a Ziploc bag. Please put the iNaturalist number (in the URL of your observations) and the species name on the voucher slips. This will save event staff a significant amount of organizational time once the collections are received.

**Figure 2.**
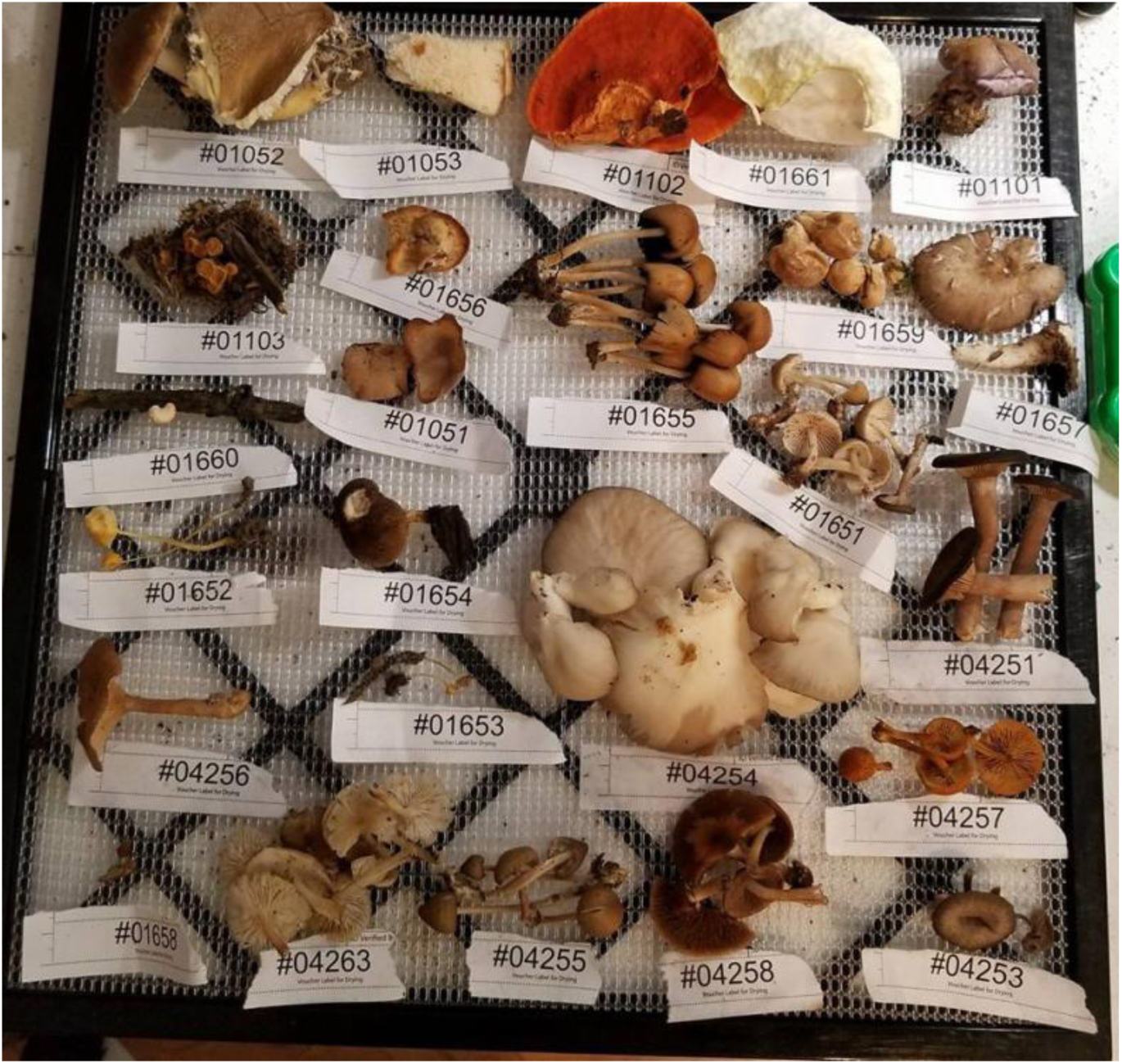
Mushroom specimens prepared for drying. Each specimen is associated with a unique number from the field data slips. These labels can be taped to the drying rack in the dehydrator.

**Figure 3.**
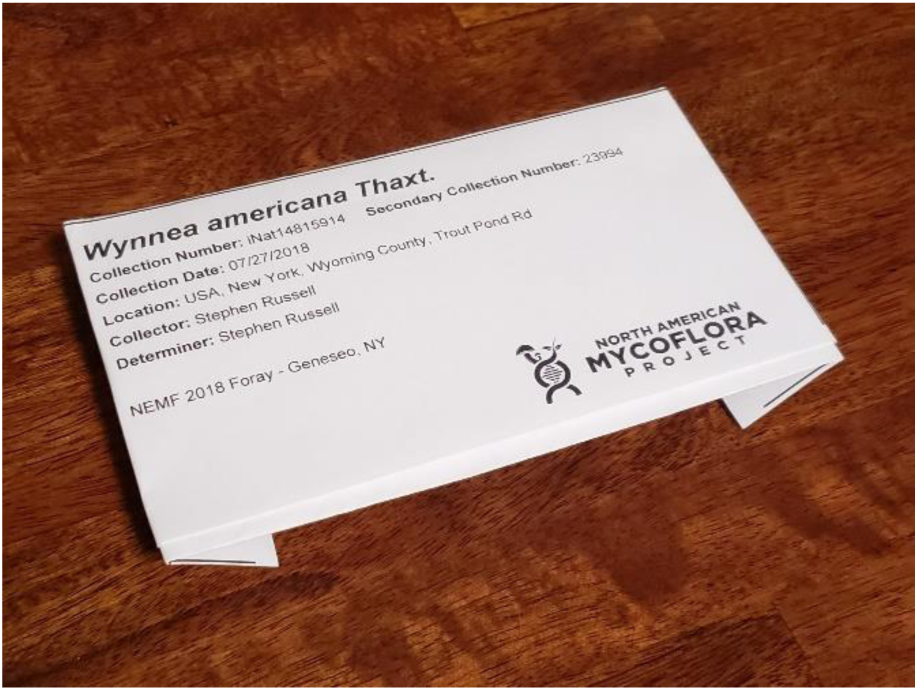
A folded specimen packet ready for submission to a herbarium.

**Figure 4.**
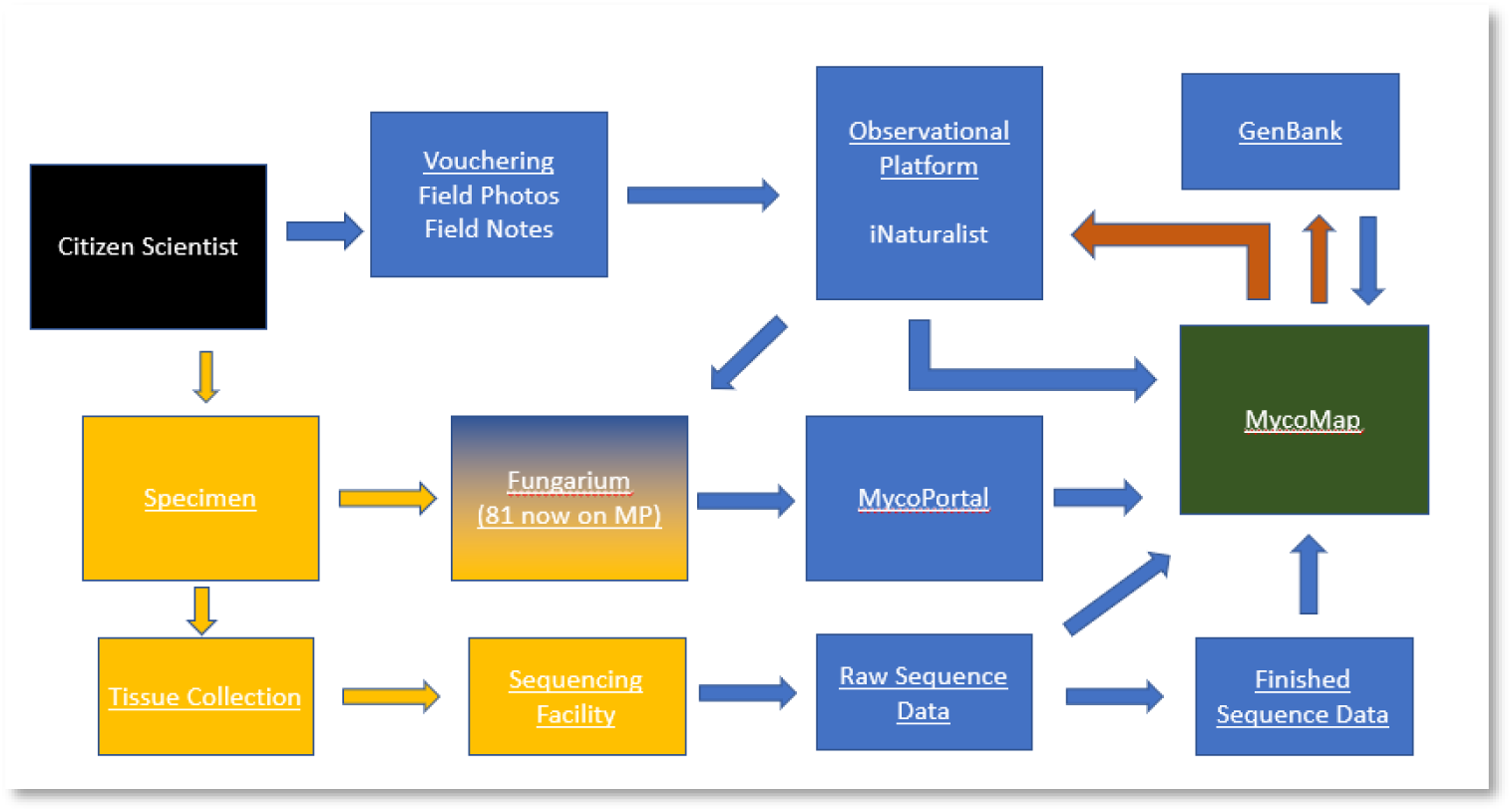
Workflow and dataflow diagram for iNaturalist online forays. Blue boxes and arrows represent data sources and data flows. Yellow boxes and arrows represent physical items and the movement of physical items. The green box represents a data aggregator. Red arrows represent data flows that have been aggregated from multiple sources.

**Figure 5.**
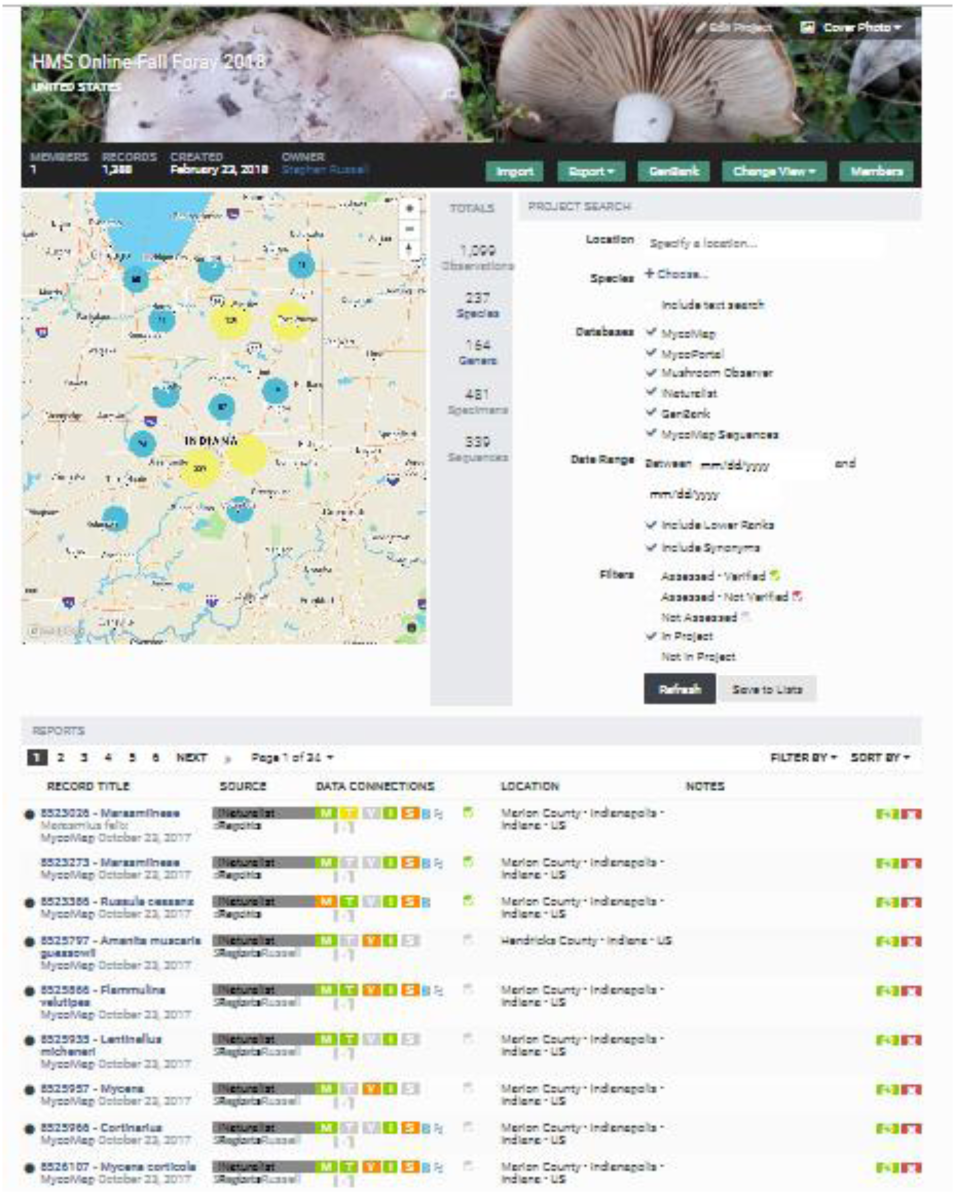
MycoMap dashboard. This interface allows for efficient management of iNaturalist project records, combined with DNA sequence data and herbarium records.

**4) Mail in your dried specimens** - Mail your specimens to the foray organizer. All of the best specimens that are collected as part of this event will have their DNA “sequenced” or examined. Multiple species that are new to science are likely to be found during this event. Your collections will make important contributions to an understanding of the fungi of Indiana.

#### Specimen shipment

Individual participants mailed their dried collections to a single, central collecting/sorting center. Participants covered the shipping cost themselves, which typically averaged between $5.00-$10.00.

#### Specimen triage/sorting

Once the specimens arrived at the sorting location, each individual collection went through a sorting process to determine which will be saved and DNA sequenced. This process was accomplished for all specimens by a single curator. Each individual collection was examined to ensure 1.) the specimen was properly dried and was in good condition (no mold, enough of the specimen was available to be useful for future research), 2.) the iNaturalist observation number matched the specimen that was received, 3.) the iNaturalist observation had the minimum required metadata, including a collection date, collection location (preferably with exact GPS coordinates), and color photographs (preferably *in situ*), and 4.) that the specimen was a valuable addition to the collecting effort. This final point depends on refined project goals and a skilled curator to make the assessment. Specimens were not retained for this study if they already had “adequate representation” from Indiana. The definition of “adequate representation” aligned with the broader Mycoflora of Indiana project goal of having sequenced specimens from multiple parts of the state, with all evidence pointing to a single species statewide (i.e. it does not appear to be a complex of multiple species nor does there appear to be cryptic species). Specimens were retained if there were a limited number of representative Indiana collections, there were a limited number of DNA sequences of the species, if the specimen could not be quickly identified to species based on macromorphology, or there were unique morphological, phenological, or ecological features present in the images of the specimen. At least one specimen from each collector was retained from each shipment. If a specimen was not selected to be permanently retained, the “Specimen Voucher Taken” observational field within iNaturalist was updated to “No – photo only.” This allowed the individual collectors to be instantly notified of the results of the sorting process. This field now has a “Not saved” option for collections that are not being retained.

#### Tissue collection

For each specimen that passed the sorting process, a small tissue sample, about the size of a grain of rice, was placed in a 1.5 mL microcentrifuge tube using a pair of forceps. The forceps were wiped with an alcohol swab and briefly flamed between each specimen. The flame was only applied long enough to burn off the alcohol. The microcentrifuge tubes were labeled with the “Tissue Sample” collection number replicate from the field data slip and sent to the lab for DNA extraction and amplification. A video of this process is available on YouTube (https://www.youtube.com/watch?time_continue=2&v=y_DI8OymzrI).

#### Herbarium storage

Once the specimens from an individual shipment were sorted, a CSV spreadsheet was downloaded from the iNaturalist project that contained the metadata for the specimens. This spreadsheet was uploaded to the MycoMap specimen packet generator (Russell 2018). This “metadata translator” generates an herbarium-approved, pre-formatted label that can be folded to permanently retain the specimen. Specimens are first wrapped in a Kimwipe and then placed in the folded packet, along with the field data slip from the collector. A video (https://www.youtube.com/watch?v=wvznFzjW83k) of this process are available. Packets were then assigned the next herbarium accession number and a secondary barcode number from the herbarium was placed on the exterior of the packet. The metadata was then entered into the herbarium’s internal database. Finally, the specimen packet was placed in the appropriate section of the herbarium collection and the metadata is uploaded to MyCoPortal (Miller and Bates 2017).

#### DNA extraction, PCR, and sequencing

DNA extractions of the selected specimens were performed using the Promega Wizard^®^ Purification Kit (Promega Corp., Madison, Wisconsin). DNA amplification focused on the nuclear ribosomal internal transcribed spacer region (nrITS), which includes the internal transcribed spacer 1, 5.8S, and internal transcribed spacer 2 regions. This is the “barcode” region for fungi and is typically 600 – 800 base pairs in length (Schoch et al. 2012). Each PCR reaction contained

12.5 μL Promega PCR Master Mix (Promega Corp., WI, USA), 9 μL water, 1.25 μL of forward and reverse primers, and 1 μL of the DNA template. The ITS1F forward primer and the ITS4 reverse primer were used for most collections (Gardes and Bruns 1993; White et al. 1990). The following PCR protocol was used: (i) initial denaturation at 95 C for 2 min, (ii) denaturation at 94 C for 1 min, (iii) annealing at 51 C for 1 min, (iv) extension at 72 C for 1 min, (v) repeat for 34 cycles starting at step 2, (vi) leave at 72 C for 2 min (vii) hold at 10C. Electrophoresis with a 1% agarose gel was used to verify successful amplification. PCR amplicons were sent to Genewiz (Genewiz, Inc., Boston, Massachusetts, USA) for sequencing. Forward and reverse strands were sequenced. The two reads were assembled using Sequencher 5.0.1 (Gene Codes Corp., Ann Arbor, Michigan, USA).

### Data management

The principal workload of foray organizers after the herbarium specimens have been filed and the DNA results have been acquired relates to the management of the significant volume of data (and data connections) that were generated as a part of the event. Hundreds of individual units of metadata needed to be transferred across multiple platforms in order to extract the full potential value of the data. The workflow diagram below outlines the high-level flows of data that occured. The MycoMap platform (www.mycomap.com) served as a laboratory information management system (LIMS) in order to track and aggregate the data for floristic projects from the most common sources. It also helped to provide a swift analytical workflow of the organized DNA barcode data and to appropriately tailor the final metadata and results for automated transfer to the required endpoints. The key data elements that needed to be crosslinked for an online foray include the source records on iNaturalist that contain the photographs and original metadata (collection date, collection location, voucher numbers, etc.), the accession information from herbaria (institution name, catalog number, etc), and the DNA data on GenBank (accession numbers). There are also several intermediate products that needed to be organized and retained, such as the raw trace files from the DNA sequencing facility and species name overrides that cannot be populated on iNaturalist. Several of the most important aspects for online foray organizers are discussed below.

#### MycoMap project creation

At the conclusion of the event, a CSV of the foray observations was downloaded from the iNaturalist project. This spreadsheet was uploaded to a new project on the MycoMap platform called “HMS Online Fall Foray 2017” (https://mycomap.com/projects/hms-fall-foray-2017). This process brings all of the iNaturalist observations from the event into the framework’s data management “dashboard” functionality.

Next, the DNA sequence data was associated with each iNaturalist observation. For the event, all filenames for raw trace files and edited DNA sequence data were formatted, organized into single ZIP files, and uploaded to the MycoMap project. Correctly formatted filenames automatically generate individual sequence records within the framework for each specimen (see mycomap.com for current formatting requirements). Each resulting sequence record has downloadable links for the raw trace files, indexes of the primers that were used for each sequence, links the sequence records to the original iNaturalist observations, and runs a local & NCBI BLAST search for each sequence that is submitted.

The next step was associating the herbarium data on MyCoPortal to the iNaturalist observations and sequences. A spreadsheet listing the MyCoPortal accession numbers (termed “occid” on MycoPortal) was uploaded to the MycoMap Project. This spreadsheet contains only two columns – the MyCoPortal accession number in a column labeled “id” and the iNaturalist observation ID in a “fieldNumber” column. The iNaturalist ID was in the format “iNat1674310.” This process links the herbarium accession information to the observational records in the MycoMap project. These three uploads (iNaturalist observation numbers, raw and edited sequence files, and MyCoPortal accession numbers) link the iNaturalist observations with the sequence data, as well as with the information about the collection at the institutional herbarium.

#### DNA sequence identity assessment

The sequence information and metadata were “verified” in preparation for upload to GenBank and submission of results back to the foray participants. First, the BLAST results were examined for each specimen. If the species name needed to be altered in light of the sequence data, it was updated on the original iNaturalist observation, rather than on the MycoMap dashboard directly. After this update, a “refresh” of the record on the MycoMap dashboard was initiated. The refresh pings the iNaturalist API, acquires the latest information about the observation, including the new species name, and updates the record in the project with the latest information.

Occasionally, due to the system utilized by iNaturalist for finalizing identifications, it is not possible to get the proper identification as the primary “name-of-record” being utilized by the observation. This typically occurs if other iNaturalist users have agreed with an incorrect name on the observation. It is often not possible to change the species name on iNaturalist unless other identifiers also change their vote for the updated name. In the cases where the species name cannot be appropriately updated on iNaturalist directly, it is updated manually on the MycoMap project dashboard by clicking the Taxonomy “T” box on the line of the target record and entering the correct species name (Figure 6) and/or utilizing the “MycoMap Species Name” observational field on iNaturalist. Provisional placeholder species names were created within the MycoMap framework for DNA sequences that could not be accurately identified to the species level based on the data in public repositories, but were likely to represent unique taxa (Ex – *Russula* “sp-IN26”).

**Figure 6.**
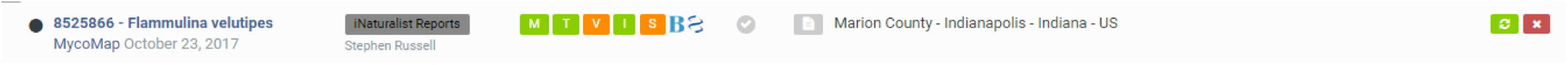
An individual line of a MycoMap project record. Users can quickly access herbarium information, raw and edited sequence data, BLAST results, and prepare sequences for upload to GenBank.

#### GenBank submission

Once the records in the MycoMap project have been corrected, the metadata for each record is approved in preparation for GenBank submission. Within the MycoMap dashboard interface, there is a NCBI icon on the dashboard line for each record that contains a DNA sequence. Clicking this icon brings up the “NCBI Verification Screen.” This screen allows individual pieces of metadata to be edited in preparation for GenBank submission (Figure 7). All fields are automatically filled out based on the metadata in the associated iNaturalist and MyCoPortal records. Once these data are verified for the sequence records, GenBank submission files can be automatically generated within the project. This includes a FASTA file containing all of the sequence data, as well as a “source modifier” TSV file that contains all of the verified metadata associated with each sequence record. These files are uploaded to the GenBank Submission Portal (https://submit.ncbi.nlm.nih.gov/subs/genbank/) to be approved for listing on NCBI’s GenBank. Once NCBI has processed and approved these records, an “Accession Report” is generated in the GenBank Submission Portal. This file (AccessionReport.tsv) is then uploaded to the MycoMap project, which links the GenBank accession numbers back to the original iNaturalist observations within the project.

**Figure 7.**
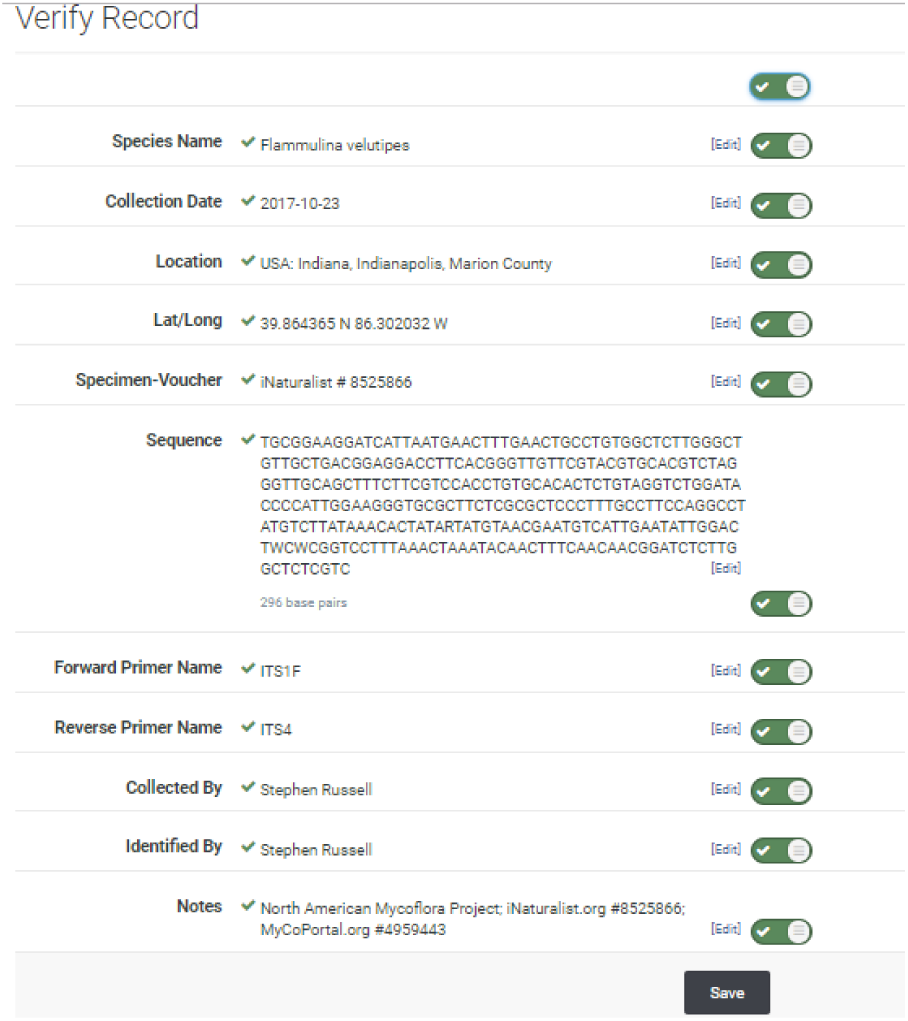
The “NCBI Verification Screen.” Metadata is formatted to NCBI specifications. Users can edit and verify individual fields in preparation for upload to GenBank.

#### Completing the circle: Pushing data back to iNaturalist observations

The final step is to publish the results of the sequence data back to the original iNaturalist observations. This notifies the foray participants that the results are ready and links the additional data that was generated back to the original iNaturalist observations. The MycoMap dashboard has a function to automatically “push” these data back to iNaturalist.

This can include links to the raw sequence files, text of the edited sequence, a link to the BLAST results, text of the GenBank accession number, text of the herbarium name and catalog number, text of the MyCoPortal accession number, and a link to the MyCoPortal record. The final iNaturalist record includes observational fields for all corresponding metadata (Figure 8).

**Figure 8.**
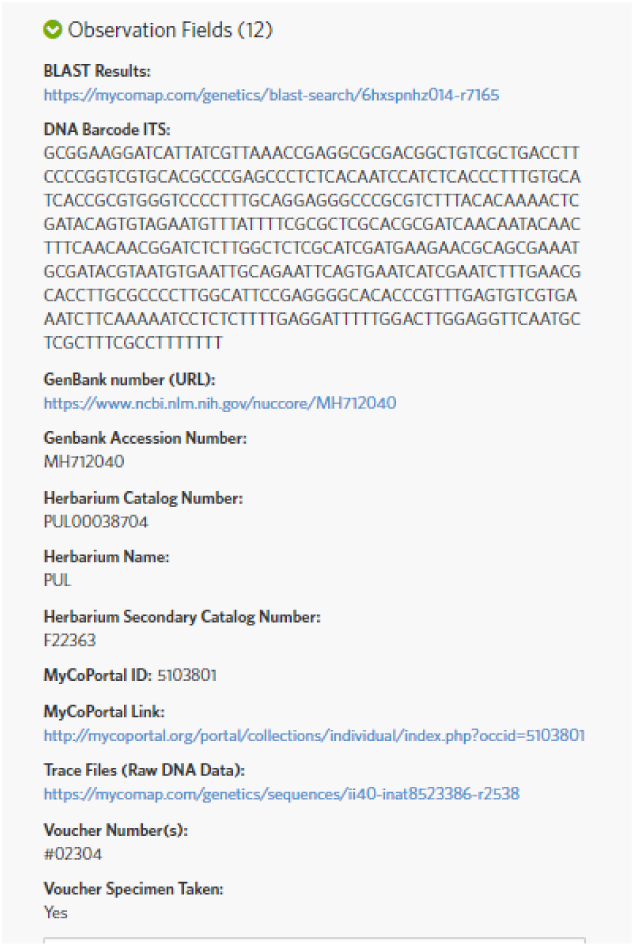
iNaturalist metadata import. An iNaturalist record with the full suite of metadata that has been automatically imported to “complete the circle” of metadata transfers.

## Results

The fall 2017 online foray resulted in 1,300+ observations from 30 participants. Over 300 species were documented, 572 specimens were stored in Purdue’s Kriebel Fungarium (PUL), and 424 of the specimens were successfully DNA sequenced. DNA extraction and amplification was attempted on all 572 of the vouchered specimens that were retained, yielding an overall sequencing success rate of 74%. DNA amplifications were only attempted a single time on these specimens. A second amplification attempt with a 1:100 dilution of the DNA template would likely increase the overall success rate. Of the 424 specimens that were successfully sequenced, 99% (420/424) could be identified to genus and 64% (273/424) could be identified to species based on reference data in public repositories (Figure 9). A total of 21% (93/424) of sequences, representing 73 putative species appear to be novel to public databases. The ten most active participants (33%) accounted for roughly 80% of the total observations (Figure 10). Table 1 outlines the species that were most frequently observed during the week of the online foray. Table 2 outlines the species which were most commonly DNA sequenced.

**Table 1.** Most frequently observed specimens from the 2017 online foray.

| Species Name | Number of Observations |
| --- | --- |
| *Trichaptum biforme* | 23 |
| *Trametes versicolor* | 22 |
| *Schizophyllum commune* | 22 |
| *Lycoperdon pyriforme* | 19 |
| *Ischnoderma resinosum* | 18 |
| *Phlebia tremellosa* | 15 |
| *Mycena haematopus* | 15 |
| *Polyporus squamosus* | 15 |
| *Pleurotus pulmonarius* | 15 |
| *Panellus stipticus* | 14 |
| *Neofavolus alveolaris* | 13 |
| *Ganoderma applanatum* | 12 |
| *Galerina marginata* | 11 |
| *Stereum ostrea* | 11 |
| *Coprinus comatus* | 10 |
| *Phellinus gilvus* | 10 |
| *Stereum complicatum* | 10 |

**Table 2.**
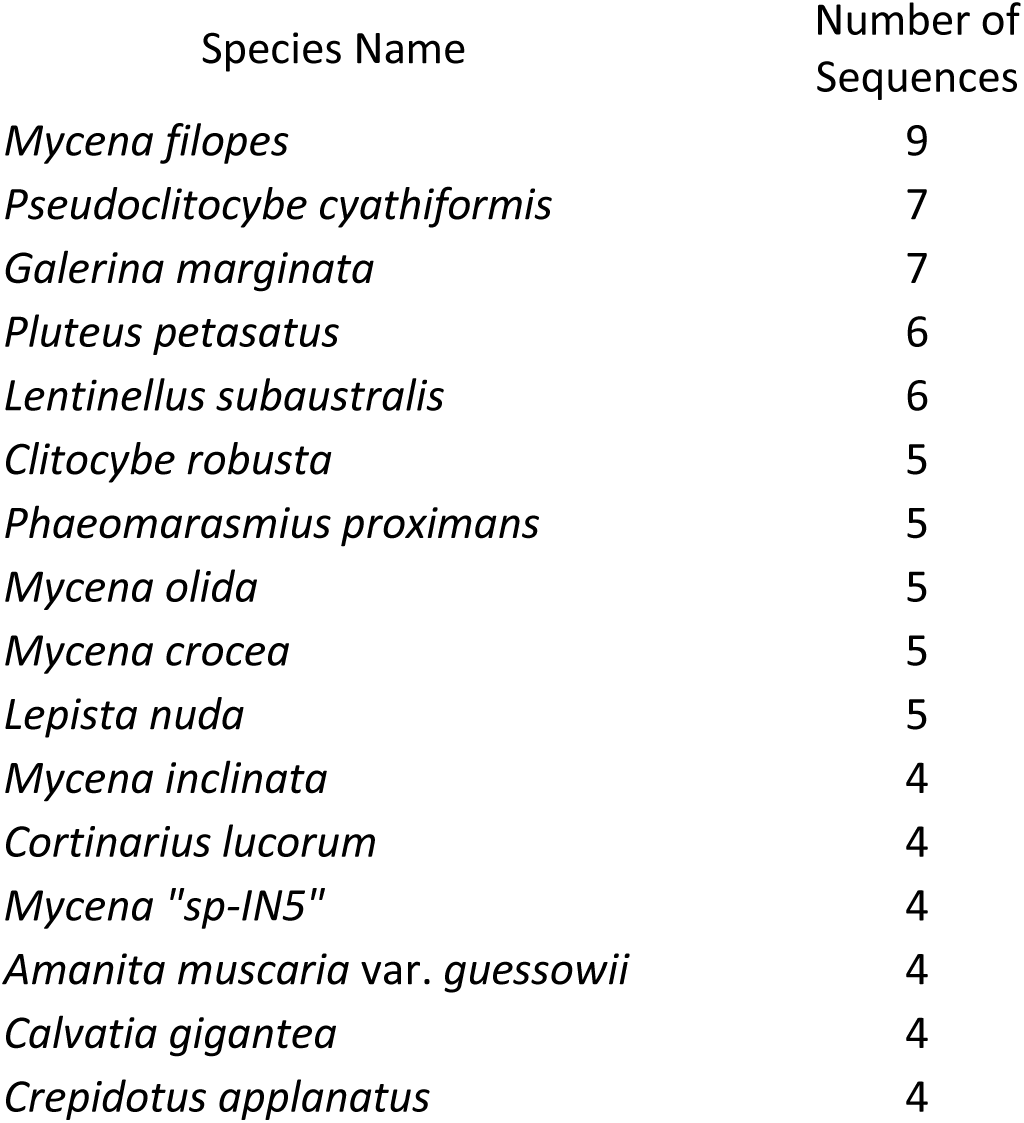
Most frequently sequenced speciess from the 2017 online foray.

| Species Name | Number of Sequences |
| --- | --- |
| *Mycena filopes* | 9 |
| *Pseudoclitocybe cyathiformis* | 7 |
| *Galerina marginata* | 7 |
| *Pluteus petasatus* | 6 |
| *Lentinellus subaustralis* | 6 |
| *Clitocybe robusta* | 5 |
| *Phaeomarasmius proximans* | 5 |
| *Mycena olida* | 5 |
| *Mycena crocea* | 5 |
| *Lepista nuda* | 5 |
| *Mycena inclinata* | 4 |
| *Cortinarius lucorum* | 4 |
| *Mycena "sp-IN5"* | 4 |
| *Amanita muscaria* var. *guessowii* | 4 |
| *Calvatia gigantea* | 4 |
| *Crepidotus applanatus* | 4 |

**Figure 9.**
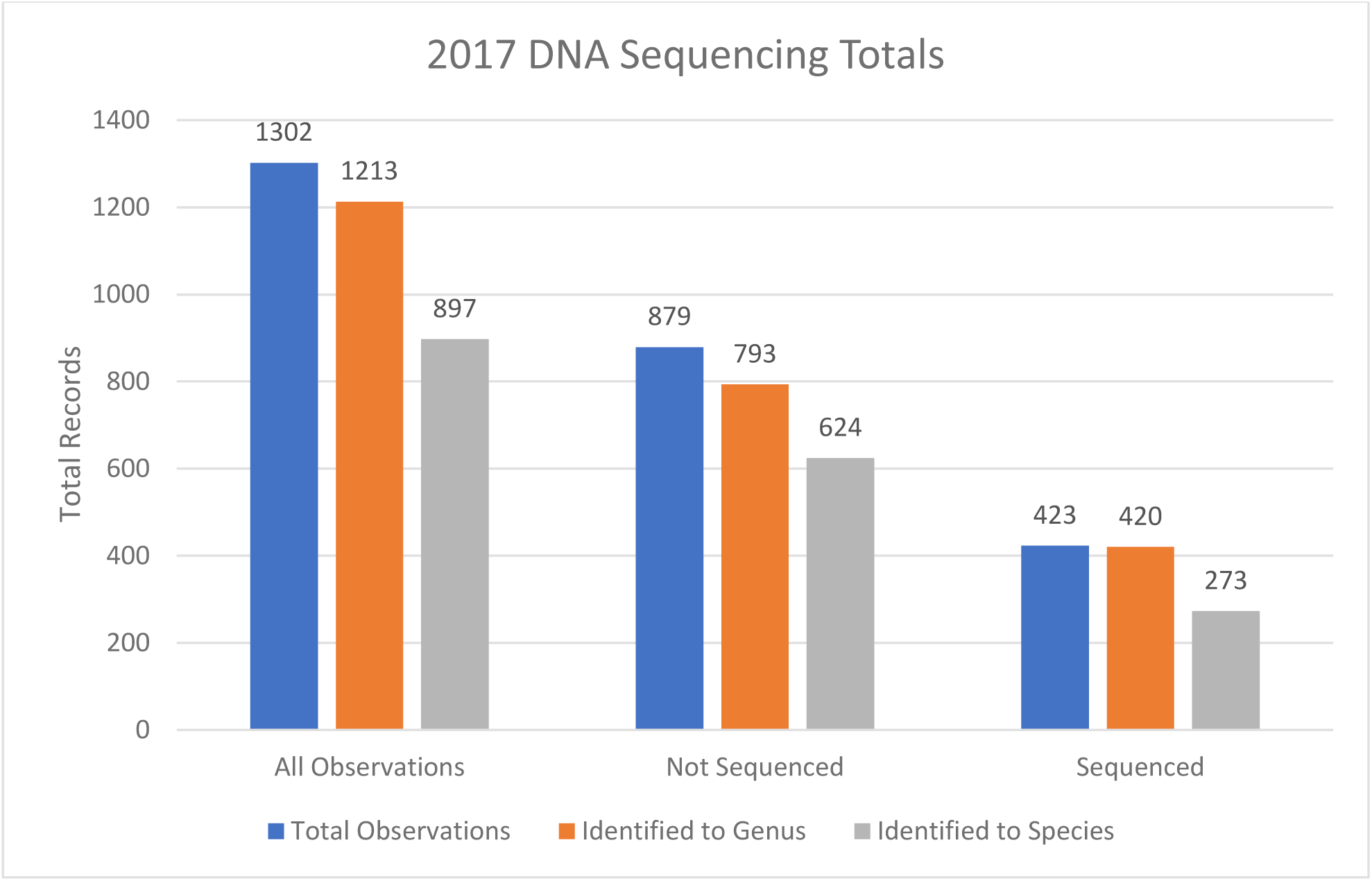
The total number of observations that were identified from the 2017 foray.

**Figure 10.**
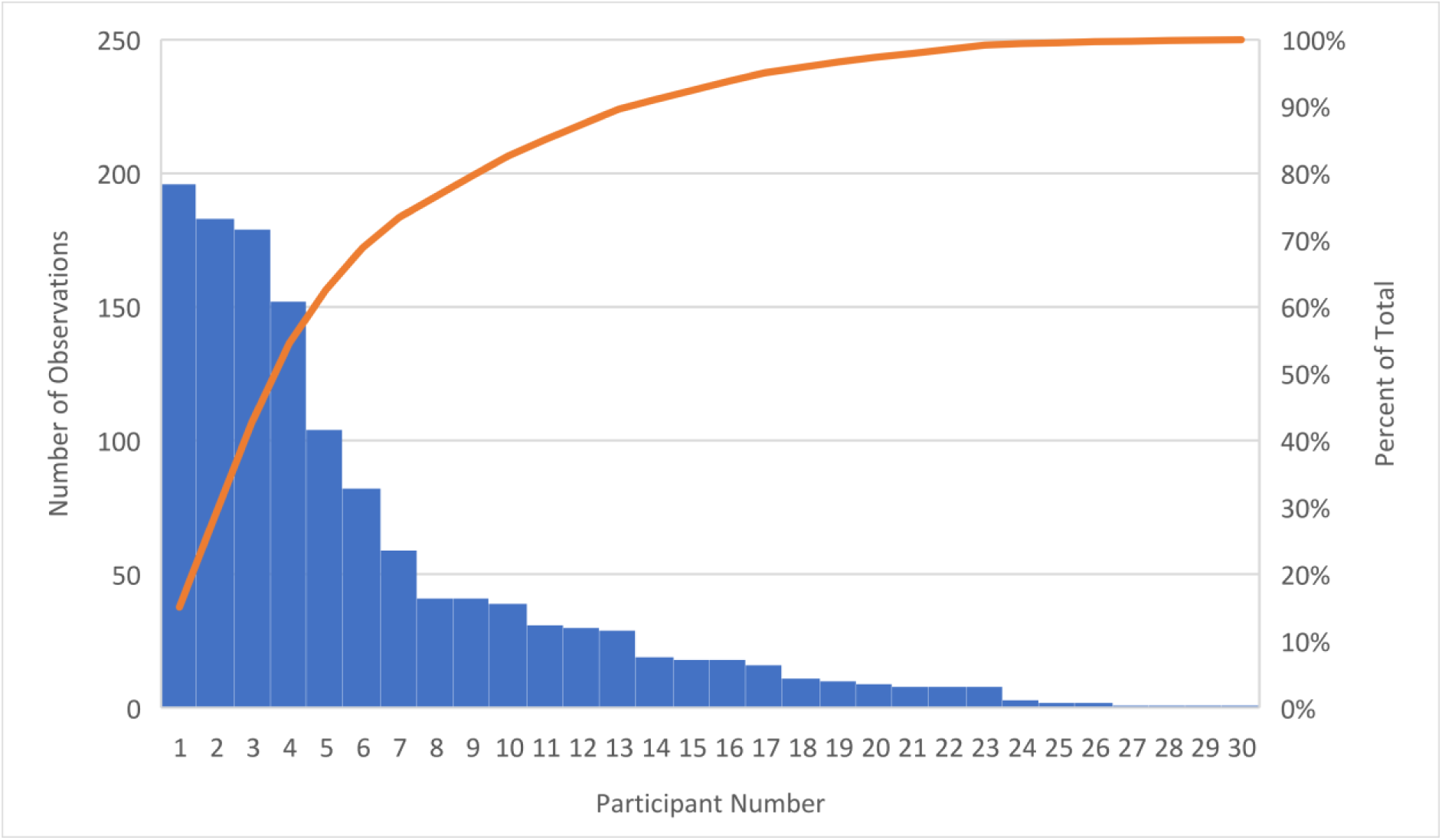
Total number of iNaturalist observations created by each participant during the 2017 foray.

## Discussion

Traditional mushroom collecting events, known as “forays,” require participants to meet at a defined geographic location where they opportunistically collect mushroom fruiting bodies as a group. For geographically diverse organizations who desire to engage citizen scientists, the traditional model presents many inherent limitations on participation. The direct costs associated with travel, potentially multiple hours each way, and the opportunity costs of the participation at a designated time (work or family obligations) present primary impediments for many citizen scientists who would otherwise be interested in participation. A traditional “bioblitz” or “mycoblitz” also suffers from many of the same limitations. The online foray outlined here has several inherent advantages. First, the event took place over the course of a week, so individuals were able to participate on their own schedule, self-selecting both the number of days they participated, as well as the duration of time they participated. Secondarily, individual collectors/participants were able to self-select collecting locations, which minimized the time and opportunity costs of participating in the survey. Participants could survey their own property or nearby publicly accessible land, rather than being required to drive to a specific location to conduct a survey as a group. This is especially important for projects that seek collections over a broad geographic area. The appearance of macrofungal fruitbodies is directly correlated with environmental conditions - primarily precipitation - and the local conditions for mushroom hunting can change significantly over short geographic ranges. Dispersing the collecting efforts over a broad geographic area allows more “prime” terrain (with the highest biodiversity of macrofungi fruiting) to be covered by collectors.

From the event organizer perspective, the online model does not take a significant amount of time or resources to plan at the front-end. A single page of information on a website was generated and an iNaturalist project was created. This process only took a couple hours. Most of the promotion for this event was conducted through the “Indiana Mushrooms” Facebook group, which has more than 25,000 members as of May 24, 2022, and directed participants to the informational website. The primary logistical issue for this inaugural event was getting field data slips to participants before the event. These field data slips, which are sequentially numbered in triplicate, proved to be an essential organizational tool, particularly because most participants had little to no experience drying and organizing physical collections. Before the event, a participant order form was placed on the website requesting participants to estimate how many specimens they planned to collect during the week of the event. Many of the users were accurate in their estimates and few additional notepads of field data slips needed to be mailed during the week of the event. For future events, sequentially numbered field data slips were available to download online and to be printed off by participants at home.

An additional purpose of these field data slips was to accurately capture size information about the specimens. Each field data slip has a centimeter scale along the top and side of the form. Participants were asked to take a picture of the field data slip with each collection, so the scale is captured with the specimen before it is dried. After drying, mushrooms lose most of their original size and shape, so recording this information early after harvest is essential if it is to be retained. Finally, the numbers allowed specimens to be sorted at the herbarium efficiently and allowed for verification that the specimens received at the centralized sorting center were assigned with the correct iNaturalist number and corresponding photographs. These types of double-checks are paramount to ensure the final data being generated maintains high fidelity throughout the process.

One organizational aspect to improve after the first event was to stress the request for participants to include the iNaturalist observation number on each field data slip. From a post-event organizational perspective, one of the activities that took the most time was to lookup the iNaturalist numbers associated with each field data slip number (and each specimen). The iNaturalist numbers were included in the final herbarium accessions for each specimen.

During the event itself, the aspect that took organizers the most time was identifying observations throughout the week on iNaturalist. Quick and regular feedback of unidentified specimens will encourage a higher level of ongoing participation. At least one pre-arranged identifier for every ten expected participants is suggested. Identifiers do not have to be locals; they can be from anywhere in the country (or world) since all of the photographs are online. There were three primary identifiers for the first online foray and these specialists identified and/or commented on the majority of the 1,300 specimens. Since the event was open online, other iNaturalist users not participating in the event also had the opportunity to provide identifications. A total of 72 individuals made identifications for the 2017 event. Had the event been much larger, more primary identifiers would have been needed.

The success of many traditional biodiversity surveys are strongly dependent upon the skill level and/or the amount of training that identifiers for the project have received. As all specimens for this project were photographed and contained a corresponding physical specimen that was DNA barcoded, neither the accuracy of the initial field identifications by participants, nor the accuracy of the online identifications by the local experts were critical to the overall project success. The primary advantage that identifiers provide during the event was to give quick initial feedback to participants about the likely identity of their collections. Eliminating the need for accurate field identification is the key aspect to this model that allows broad participation of members from every skill level, without the need for extensive training before they can make effective contributions to the project.

Despite few participants having experience with saving/drying specimens, the vast majority of the specimens that came in were of good quality and were exceptionally well-organized. The top collectors for this event provided over 150 high quality specimens, yet had not previously participated in any large-scale efforts to save macrofungal collections. Furthermore, the only training they received was the short set of protocols described above. During the 2017 event, only one set of approximately 20 specimens, from a single individual, failed to follow the organizational protocols.

It is possible that this model will not work equally well in all areas, as many states and local governmental units have strict regulations regarding harvesting mushrooms in public areas. Organizers need to ensure their participants are aware of local restrictions on mushroom collection to confirm they are complying with all local laws and do not risk citations. Most regulatory bodies have a system of granting scientific collecting permits, so obtaining permission for participants to access otherwise restricted areas may be possible. However, this process typically takes a substantial amount of time for authorization, so the planning process would need to extend well before the event takes place.

One possible point of failure for these events is that participants do not end up mailing in the specimens they collected. Despite participants being required to cover shipping costs themselves, nearly all of the specimens that were flagged as containing a specimen were actually received at the central collecting center. Drying and organizing specimens takes a significant amount of time, so there are built-in incentives for users to actually complete the process so their work is not for naught. An additional built-in incentive is that a many participants desire to have their specimens DNA sequenced so they can get granular feedback as to how their personal collections may be contributing directly to the knowledge of biodiversity of the state. This type of post-event feedback is essential for future participation and can be easily accomplished by posting results back onto the iNaturalist platform. The MycoMap dashboard streamlines this feedback process. The prizes that were offered did not seem to be an essential component for participation, but it did add a “gamification” aspect to the event that likely increased the competitive drive for some of the top-performing participants, and added another layer of interest to the overall design of the event.

The post-event resources that are available should be one of the top considerations for event organizers. With the model outlined here, foray organizers were willing to accept and sort through all of the specimens that were collected at the event. This typically involved 500-800 specimens. It takes a significant amount of time to organize, verify metadata, print labels, and collect tissue for this many specimens. Even if there is the workforce available to perform these tasks, there must be an institutional herbarium that is willing to accept the specimens. A principal reason that foray organizers desired to sort all of the specimens is that it is preferable for experts to make the determination if an individual specimen is important enough to save, rather than untrained citizen scientists making the determination. For example, there would likely be a deficit of small, boring brown mushrooms if citizen scientists were to make the determination to retain a specimen or not, along with an overrepresentation of larger, more colorful species. Typically, the larger and more colorful a specimen is, the higher the likelihood the species has already been thoroughly described in the literature and has been previously DNA sequenced. If the ability to accept a large number of specimens is a limiting factor, a different method for specimen collection may be required. Other options would include focusing citizen scientists on a small number of target genera or having skilled identifiers request individual specimens during the event, instead of having participants collect, organize, and dry the majority of the specimens they encounter.

Finally, the amount of funding available for DNA barcoding is also likely to be a limiting factor for anyone planning a similar event. DNA sequencing costs represented the highest direct cost. Labor was donated for the DNA extraction, amplification, and sequencing preparation for of all of the specimens at this event, so the total cost of DNA barcoding was under $5.00 per specimen. This meant the required budget to sequence 500 collections per event was under $2,500 – well within the budget for many organized mycological societies across North America. Without individual relationships at a partner molecular lab, the costs can increase to $20+ per specimen. This would significantly increase the required budget for an event at this scale, or it would require that foray organizers have to design far more stringent protocols on which specimens are retained for DNA sequencing.

## Acknowledgements

The authors thank members of the Hoosier Mushroom Society who participated in this event. The time and dedication by these members are what has made these events possible and scientifically valuable. Special thanks also goes to the Dr. Aime and the Aime Lab at Purdue University for providing the laboratory space and equipment that was utilized for DNA barcoding of the specimens, as well as to the graduate and undergraduate students of the lab who participated in this event. Thanks to Purdue’s Kriebel Fungarium for accepting the specimens that were submitted. Thank you to Maggie Russell, who gave commentary on the final manuscript. Finally, thanks to the undergraduates of Dr. Peter Avis at Indiana University Northwest and of Dr. Scott Bates at Purdue University Northwest for contributing collections to these events.

## Competing Interests

The author(s) has/have no competing interests to declare.

## Notes

### Competing Interest Statement

The authors have declared no competing interest.

## References

Bates, S.T., Miller, A.N. and the Macrofungi Collections and Microfungi Collections Consortia, 2018. The protochecklist of North American nonlichenized Fungi. Mycologia, 110(6), pp.1222–1348. DOI: https://doi.org/10.1080/00275514.2018.1515410

Blackwell, M., 2011. The Fungi: 1, 2, 3… 5.1 million species?. American journal of botany, 98(3), pp.426-438. DOI: https://doi.org/10.3732/ajb.1000298

Bruns, T.D., and M.W. Beug. 2012. Working toward a North American Mycoflora for Macrofungi. McIlvainea 21. Available at http://www.namyco.org/working_toward_a_north_america.php

Buyck, B., Olariaga, I., Justice, J., Lewis, D. and Hofstetter, V., 2016. The dilemma of species recognition in the field when sequence data are not in phase with phenotypic variability. Cryptogamie, Mycologie, 37(3), pp.367-390. DOI: https://doi.org/10.7872/crym/v37.iss3.2016.367

Hawksworth, D.L. and Lücking, R., 2017. Fungal diversity revisited: 2.2 to 3.8 million species. The fungal kingdom, pp.79-95. DOI: https://doi.org/10.1128/9781555819583.ch4

Hawksworth, D.L. and Rossman, A.Y., 1997. Where are all the undescribed fungi?. Phytopathology, 87(9), pp.888–891. DOI: https://doi.org/10.1094/PHYTO.1997.87.9.888

Korhonen, A., Seelan, J.S.S. and Miettinen, O., 2018. Cryptic species diversity in polypores: the Skeletocutis nivea species complex. MycoKeys, (36), p.45. DOI: https://dx.doi.org/10.3897%2Fmycokeys.36.27002

Korf, R.P., 2005. Reinventing taxonomy: a curmudgeon’s view of 250 years of fungal taxonomy, the crisis in biodiversity, and the pitfalls of the phylogenetic age. Mycotaxon, 93, pp.407–416.

Miller, A.N. and Bates, S.T., 2017. The Mycology Collections Portal (MyCoPortal). IMA Fungus, 8(2), pp.65-66. DOI: https://doi:10.1007/BF03449464

Pimm, S.L., Jenkins, C.N., Abell, R., Brooks, T.M., Gittleman, J.L., Joppa, L.N., Raven, P.H., Roberts, C.M. and Sexton, J.O., 2014. The biodiversity of species and their rates of extinction, distribution, and protection. Science, 344(6187), p.1246752. DOI: https://10.1126/science.1246752

Russell, S.D. 2018. Specimen Packets. North American Mycoflora Project. Available at: http://mycoflora.org/component/sppagebuilder/59-specimen-packets [Last accessed 30 December 2018]

Schoch, C.L., Seifert, K.A., Huhndorf, S., Robert, V., Spouge, J.L., Levesque, C.A., Chen, W. and Fungal Barcoding Consortium, 2012. Nuclear ribosomal internal transcribed spacer (ITS) region as a universal DNA barcode marker for Fungi. Proceedings of the National Academy of Sciences, 109(16), pp.6241-6246. DOI: https://doi.org/10.1073/pnas.1117018109

Seltzer, C. 2018. Announcing Changes to Projects on iNaturalist. Available at: https://www.inaturalist.org/blog/15450-announcing-changes-to-projects-on-inaturalist [Last accessed 30 December 2018]

Sheehan, B., 2017. Mushroom citizen science in the USA: From species lists to Mycoflora 2.0. Fungi Magazine, 101, pp.28–36.

Seifert, K.A., 2009. Progress towards DNA barcoding of fungi. Molecular ecology resources, 9, pp.83-89. DOI: https://doi.org/10.1111/j.1755-0998.2009.02635.x

Truong, C., Mujic, A.B., Healy, R., Kuhar, F., Furci, G., Torres, D., Niskanen, T., Sandoval-Leiva, P.A., Fernández, N., Escobar, J.M. and Moretto, A., 2017. How to know the fungi: combining field inventories and DNA-barcoding to document fungal diversity. New Phytologist, 214(3), pp.913-919. DOI: https://doi.org/10.1111/nph.14509

United States Postal Service 2018. Domestic Mail Manual: 483 Rates and Eligibility. Available at: https://pe.usps.com/archive/html/dmmarchive20070315/483.htm [Last accessed 30 December 2018]

Yahr, R., Schoch, C.L. and Dentinger, B.T., 2016. Scaling up discovery of hidden diversity in fungi: impacts of barcoding approaches. Philosophical Transactions of the Royal Society B: Biological Sciences, 371(1702), p.20150336. DOI: https://doi.org/10.1098/rstb.2015.0336

